# Gains in soil C storage under anthropogenic N deposition are rapidly lost following its cessation

**DOI:** 10.1101/2023.10.31.562515

**Authors:** Brooke E. Propson, Donald R. Zak, Aimée T. Classen, Andrew J. Burton, Zachary B. Freedman

## Abstract

Anthropogenic nitrogen (N) deposition enhanced the global terrestrial carbon (C) sink and partially offset anthropogenic CO_2_ emissions. This was driven by the suppression of microbial activity associated with the breakdown of soil organic matter. However, since the implementation of emission abatement policies in the 1970’s, atmospheric N deposition has declined globally, and the consequences of this decline are unknown. Here, we assessed the response of soil C storage, and associated microbial activities, in a long-term field study that experimentally increased N deposition for 24-years. We measured soil C and N, parameters of microbial activity, and compared effect sizes of soil C in response to, and in recovery from, the N deposition treatment across the history of our experiment (1994-2022). Our results demonstrate that the accumulated C in the organic horizon has been lost and exhibit additional deficits 5-years post-termination of the N deposition treatment. These findings, in part, arise from mechanistic changes in microbial activity. Soil C in the mineral soil was less responsive thus far in recovery. If these organic horizon C dynamics are similar in other temperate forests, the Northern Hemisphere C sink will be reduced and climate warming will be enhanced.

## INTRODUCTION

Anthropogenic nitrogen (N) deposition in Northern Hemisphere temperate forests has increased forest soil carbon (C) storage, thereby slowing atmospheric CO_2_ accumulation and subsequent climate warming^1–3^. To better understand the mechanisms mediating the ecosystem-level response to N deposition, we have conducted a 36-year field experiment to simulate future rates of anthropogenic N deposition. To date, experimental N deposition, herein referred to as the N deposition treatment, substantially increased soil organic matter (SOM) accumulation in the forest floor (+51%) and the mineral soil (+18%)^4^. This increase in soil C is attributable to mechanistic changes in microbial function, namely the suppression of microbial activity associated with the breakdown of biochemically recalcitrant SOM (Supplementary Table 1). This results in reduced rates of leaf litter and SOM decay and the resulting accumulation of C in soil^3–12^. These findings suggest a potential mechanism supporting the extensive, unresolved terrestrial C sink in the Northern Hemisphere, accounting for 15-30% of annual global anthropogenic emissions^13,14^. Further, evidence suggests that anthropogenic N deposition can increase soil C sequestration across a wide array of terrestrial ecosystems, regardless of ecosystem N status and climate zones^8,15-17^.

Subsequent to emission abatement policies developed under the United Nation’s Convention on Long-range Transboundary Air Pollution (CLRTAP) in 1979, global anthropogenic emissions and deposition of atmospheric N have declined since the 1990’s^18–22^. In temperate forests of the Upper Great Lakes region, for example, wet atmospheric NO_3_ deposition in the 1980’s averaged between 8.5-18.5 kg NO_3_ ha^-1^yr^-1^ and was projected to reach 30 kg NO_3_ ha^-1^yr^-1^ by 2050. However, as of 2021, wet atmospheric NO_3_ deposition in the region averaged between 5-8 kg NO_3_ ha^-1^yr^-1^ (https://nadp.slh.wisc.edu/). It is plausible that reduced N deposition, as legislated under the CLRTAP, may reduce the soil C sink supported by northern hardwood forests and may ultimately feedback to enhance climate warming. Thus, it is imperative to understand how forest ecosystems, and the terrestrial C sink that they support, recover from historically high rates of atmospheric N.

Conceptual models hypothesize that ecosystem response to reduced N deposition will exhibit a hysteretic model of recovery, in which the time from when atmospheric N deposition is reduced and ecosystem-level changes are detectable (i.e., lag-time) will be different for different ecosystem pools and fluxes^23,24^. Here, we assess the initial recovery of the terrestrial C sink that was supported under high rates of anthropogenic N deposition. Building upon the hysteretic model of recovery proposed by Gilliam et al. (Ref. 23), and adapted by Carrara et al. (Ref. 25), we broadly hypothesize that the terrestrial C sink supported by high rates of anthropogenic N deposition will exist across a spectrum of recovery in various states (i.e., conditions). Specifically, the soil C pool may exist as either retained in the system in the high N deposition state (condition 1), returned to the ambient state (condition 2), or shifted towards a new steady state (condition 3) (Fig. 1). Sugar maple leaf litter decays rapidly and thus, we hypothesize that organic horizon C will be depleted during the initial recovery and return to the ambient state (condition 2; Fig. 1; H1a)^4,26^. We hypothesize that the accumulated mineral soil C in the N deposition treatment will persist and thus changes in mineral soil C will not be observed in initial recovery (condition 1; Fig. 1; H1b)^23^. Finally, we hypothesize that microbial activity in the organic and mineral horizons will parallel our predicted soil C responses, with microbial activity returning to that of the ambient condition in the organic horizon (condition 2; Fig. 1; H2a) and remaining suppressed in the mineral soil (condition 1; Fig. 1; H2b). To address these alternatives, we utilized a long-term field study that received experimentally increased N deposition for 24-years in MI, USA (Fig. 2) and examined the response of C pools and soil microorganisms 5-years post-termination of the N deposition treatment.

**Figure 1.**
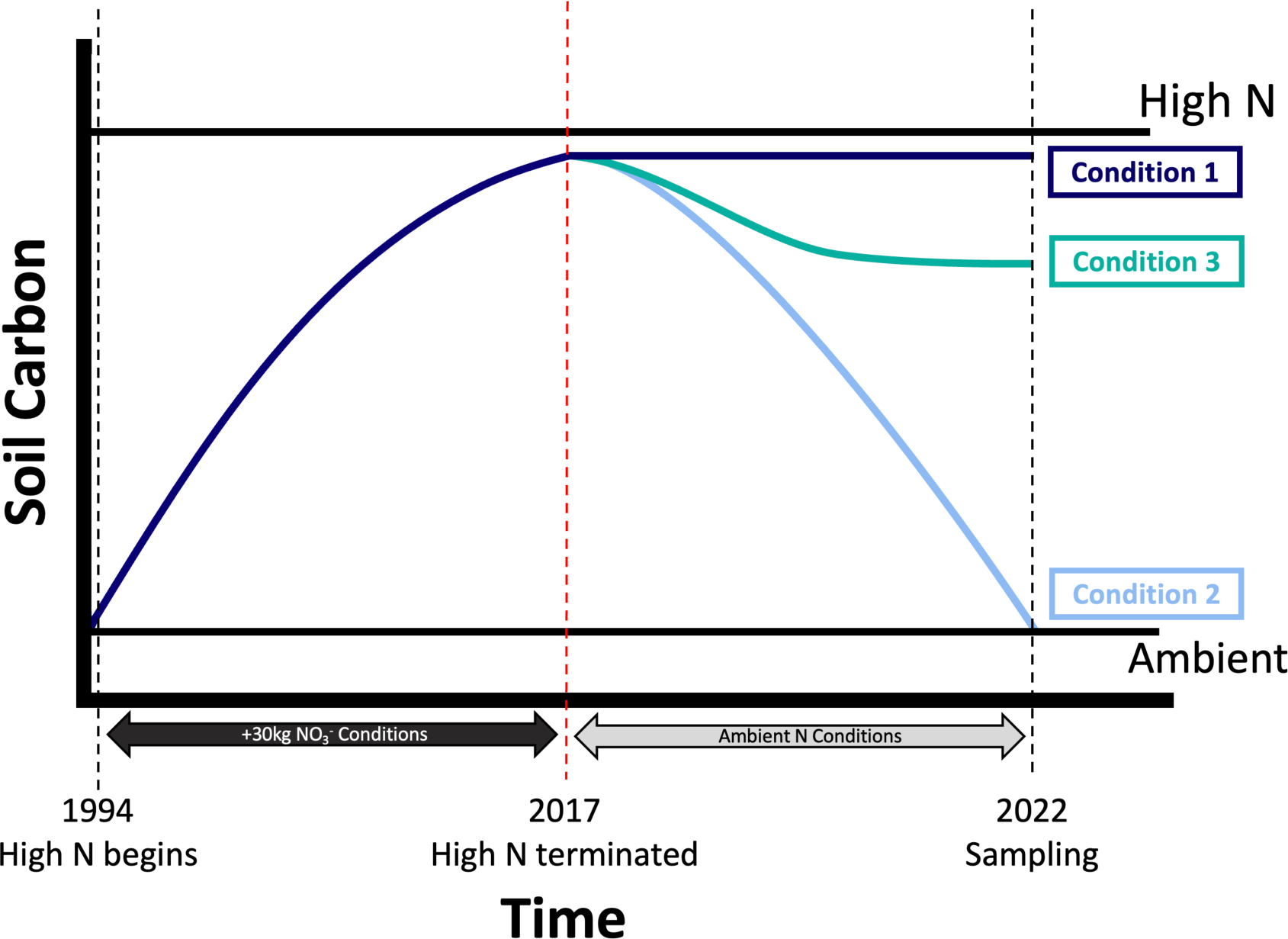
Proposed conceptual model of recovery from historically high N deposition. Soil C that accumulated under high N deposition will exist across a spectrum of recovery as either retained in the system in the high N deposition state (condition 1), returned to the ambient state (condition 2), or have begun to shift to a new steady state (condition 3).

**Figure 2.**
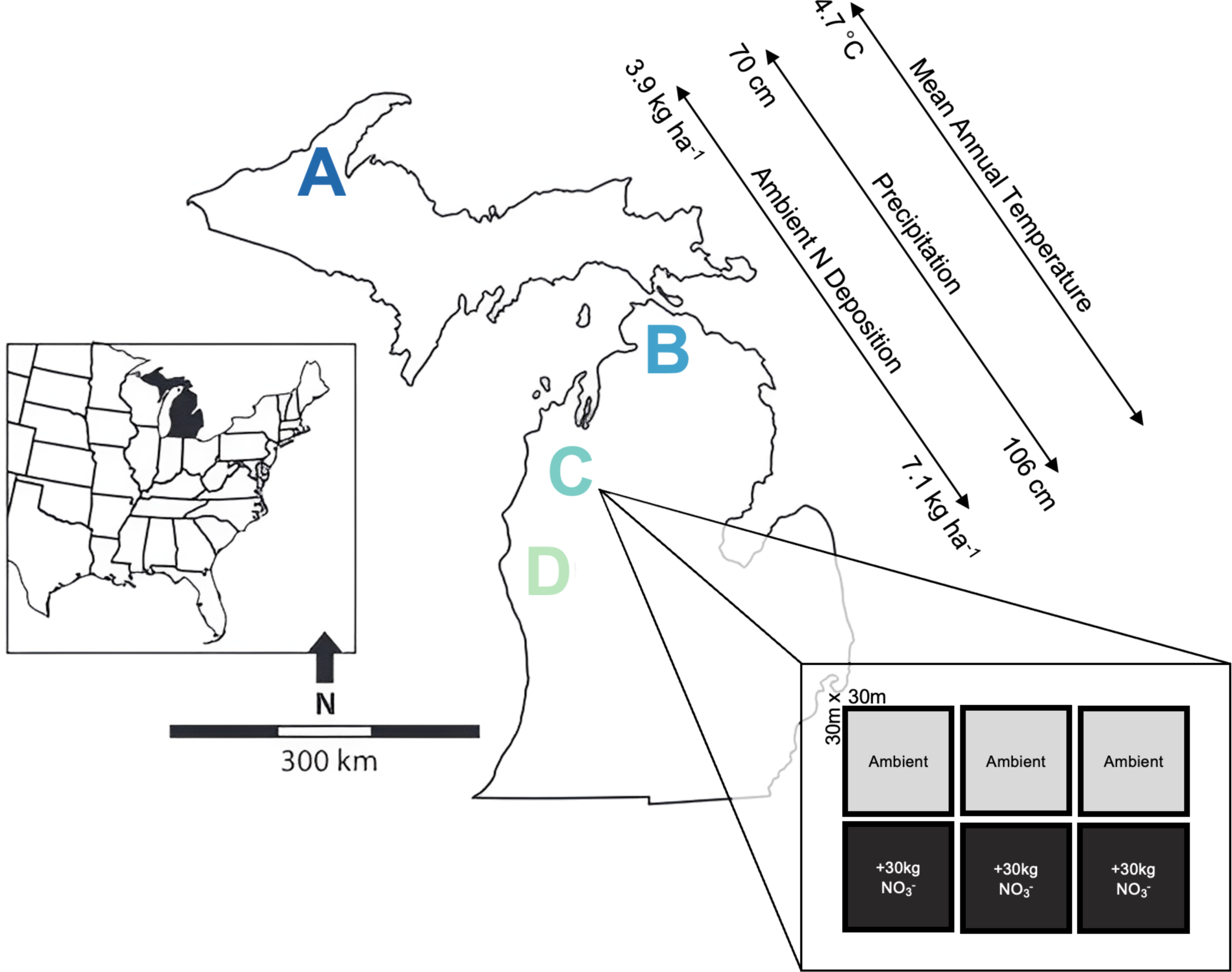
The distribution of study sites in Michigan, USA that span the geographic range of northern hardwood forests. The sites were selected due to their similarity of floristic and edaphic characteristics (see Supplementary Table 2). At each site (n=4), there are six 30×30-m plots, three of which received experimental N additions in the form of 30 kg NO_3_^—^N ha^-1^ yr^-1^ from 1994-2017 and three of which receive ambient atmospheric N deposition.

## RESULTS

### Soil C Storage

Five years post-termination of the experimental N deposition treatment, the previously documented increases in organic horizon mass (+51% change from the ambient condition; Supplementary Table 1) and SOM content (+18%; Supplementary Table 1) in the N deposition treatment were no longer present (Fig. 3). While this pattern of reduced organic horizon C was strong at the three most southern sites (i.e., sites B, C, and D), the northernmost site A still exhibited greater organic horizon mass (+73%, *P* ≤ 0.001; Extended Data Fig. 1) and soil C content (+77%, *P* ≤ 0.001; Extended Data Fig. 2) in the N deposition treatment relative to ambient conditions, despite not receiving experimental N additions for 5 years. Further, at the three southernmost sites (sites B, C, and D), the organic horizon C was significantly reduced in the N deposition treatment as compared to organic horizon C observed under ambient conditions. Specifically, in the N deposition treatment at sites B, C, and D, we observed −44%, −48%, and −41% reductions in soil organic horizon C, respectively (all *P* ≤0.001; Extended Data Fig. 2), compared to ambient conditions 5-years post-termination of the N deposition treatment.

**Figure 3.**
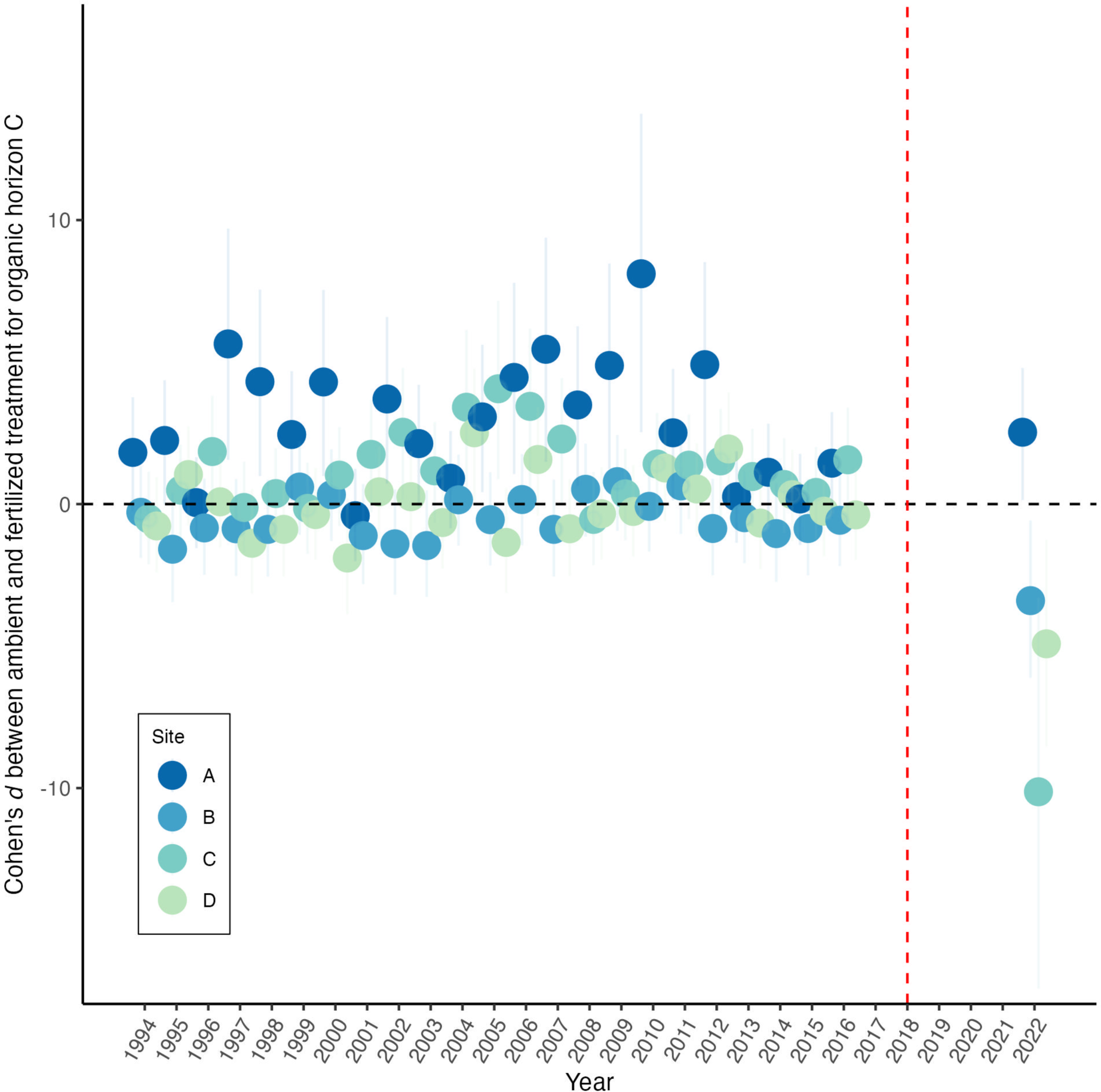
Standardized effect sizes (Cohen’s d) with 95% confidence intervals of the N deposition treatment on organic horizon C throughout the long-term experiment. The N deposition treatment was applied annually from 1994-2017. The red vertical line indicates the first year in which no treatment was applied. Points are colored by site (n = 4). Points above zero indicate more organic horizon C in the N deposition treatment compared to the ambient treatment. Points below zero indicate more organic horizon C in the ambient treatment compared to the N deposition treatment.

Across all sites, the increased mineral soil C that accumulated under experimental N deposition remained observable 5-years post-termination of the N deposition treatment (Fig. 4a), though the increase (+11%) is no longer statistically significant *P* = 0.31; Fig. 4b). Site and treatment did not significantly interact to influence mineral soil C content.

**Figure 4.**
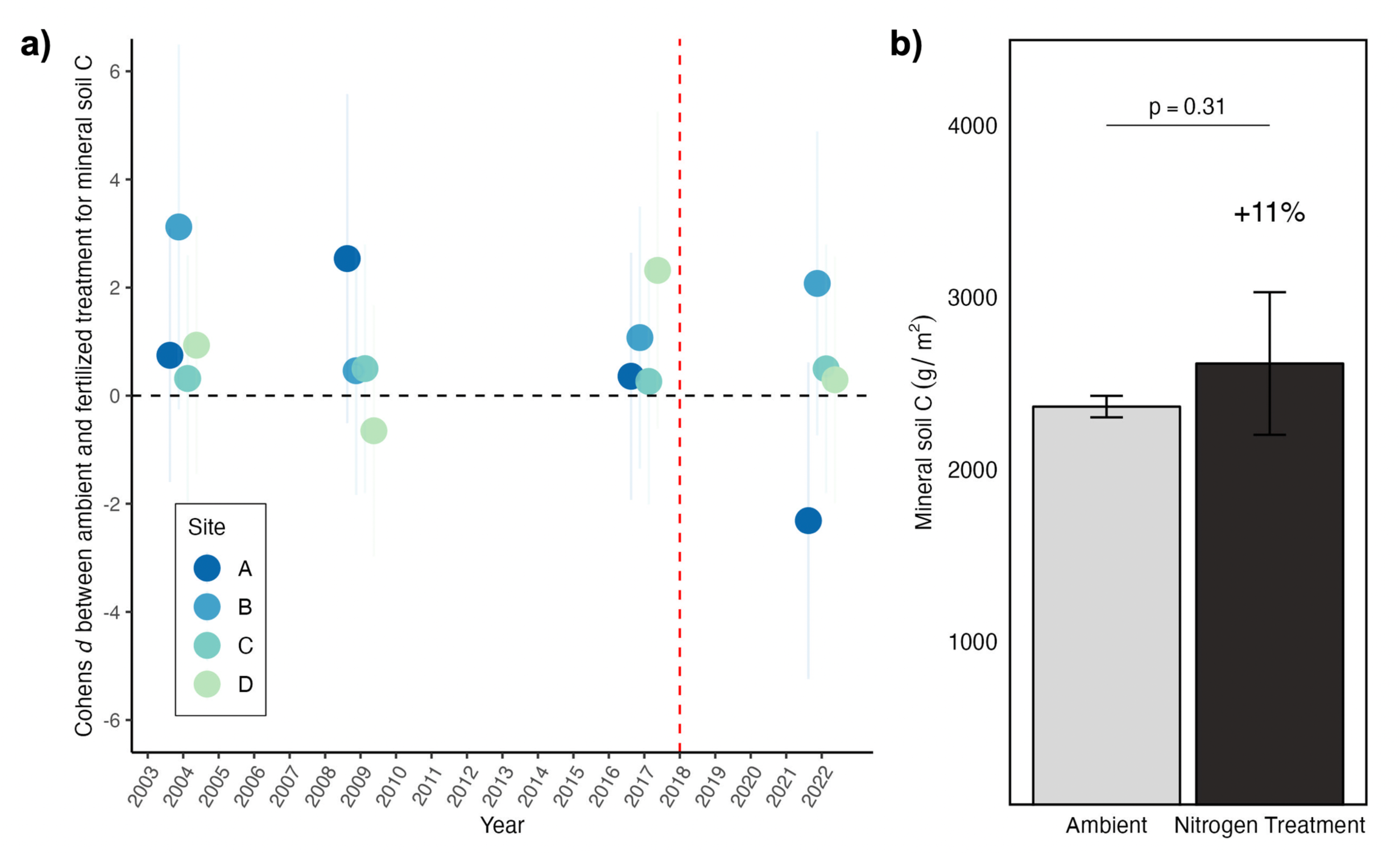
Standardized effect sizes (Cohen’s d) with 95% confidence intervals of the N deposition treatment on mineral soil C throughout the long-term experiment. The N deposition treatment was applied annually from 1994-2017. The red vertical line indicates the first year in which no treatment was applied. Points are colored by site (n = 4). Points above zero indicate more mineral soil C in the N deposition treatment compared to the ambient treatment. Points below zero indicate more mineral soil C in the ambient treatment compared to the N deposition treatment **(a)**. The effect of the historically high N deposition on mineral soil C values from samples collected in September 2022, 5-years post-termination of the N deposition treatment. Values represent treatment means ± standard error (n=12). Site × Treatment *P* > 0.05 **(b)**.

### Microbial Activity

Across all sites, there was no difference in microbial respiration between the ambient and experimental N deposition treatment in the organic horizon (*P* = 0.40) or mineral soil (*P* = 0.52) C pools and there were no significant interactions between site and treatment.

The previously documented decrease in organic horizon peroxidase (PER) activity (−50%) in the N deposition treatment was no longer observed and instead exhibited a +22% increase compared to ambient conditions across study sites (*P* = 0.03; Fig. 5). Organic horizon polyphenol oxidase (PPO) activity potential (previously −35% in N deposition treatment) was no longer suppressed in sites A, C, and D, yet remained suppressed in site B in the N deposition treatment (−43%; Site × Treatment *P* = 0.02; post hoc *P* = 0.03; Extended Data Fig. 3). Also no longer present is the previously documented decrease in organic horizon β-glucosidase (BG) activity potential (−24%) in the experimental N deposition treatment. No difference in BG activity was observed between ambient conditions and the N deposition treatment (*P* = 0.74).

**Figure 5.**
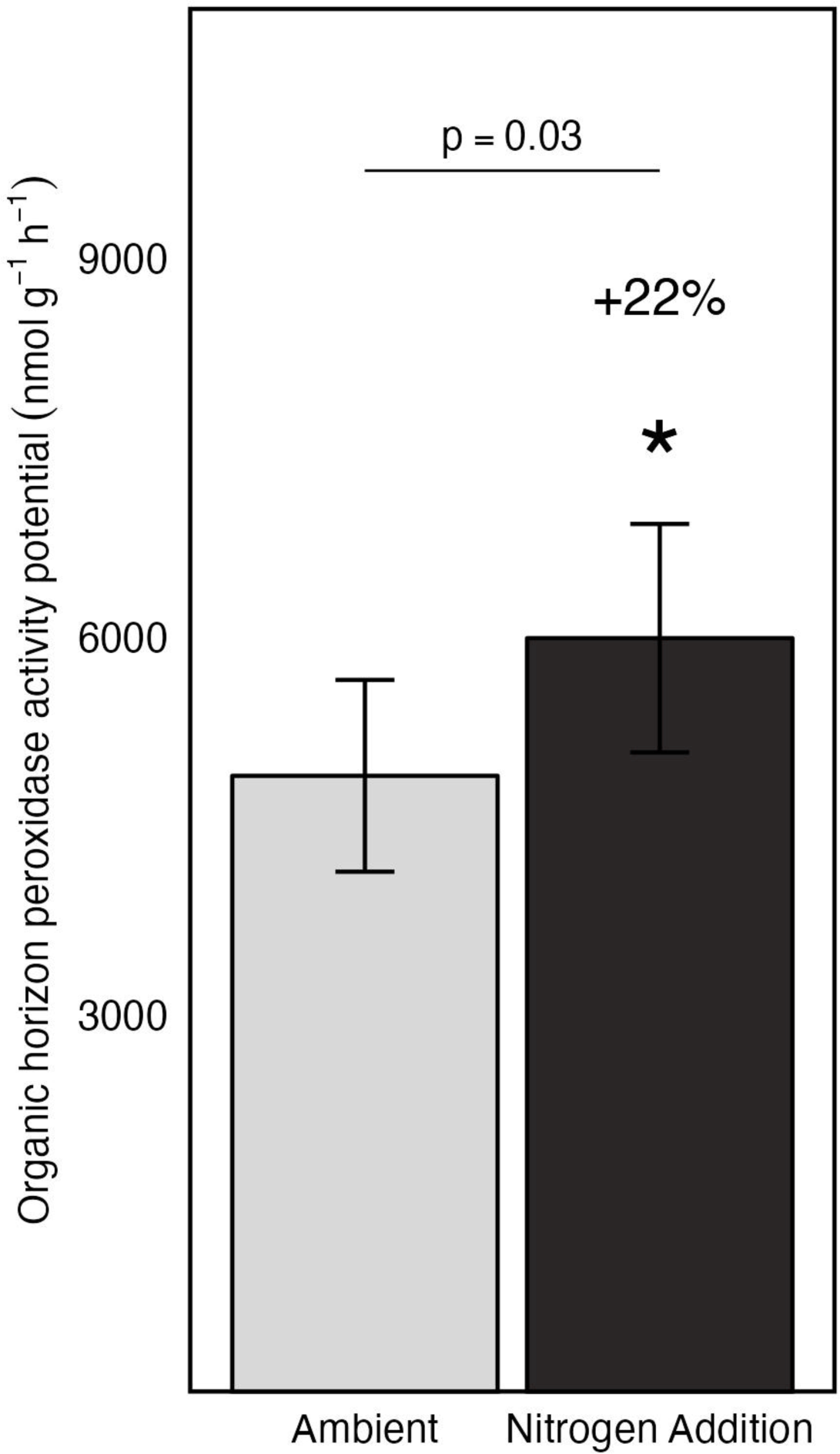
The effect of historically high N deposition on organic horizon peroxidase (PER) activity values from samples collected in September 2022, 5-years post-termination of the N deposition treatment. ANOVA was performed on estimated marginal means, but raw values are presented. Values represent treatment means ± standard error (n=12). Site × Treatment *P* > 0.05. Organic horizon PER values in the N deposition treatment percent change from ambient is reported.

In the mineral soil, the previously documented suppression of extracellular enzyme activity in the experimental N deposition treatment marginally persists, but has largely returned to levels similar to ambient conditions. Though not statistically significant, mineral soil PER activity potential in the N deposition treatment exhibited a −18% decrease relative to ambient conditions (*P* = 0.06; Extended Data Fig. 4). The previously documented suppression in mineral soil PPO and BG activity potential in the N deposition treatment is no longer present across sites (*P* = 0.99; *P* = 0.25, respectively).

## DISCUSSION

Overall, the results of this study indicate that the global terrestrial C sink in the northern hemisphere, which offset 15-30% of anthropogenic CO_2_ emissions in Earth’s atmosphere^13^, could decline due to reduction in atmospheric N deposition, potentially enhancing climate warming. We show that the increased soil C pool that accumulated under experimental N deposition was not observable 5-years post-termination of the N deposition treatment, and this pattern is, in part, the result of mechanistic changes in microbial activity. This is the first report of forest recovery from long-term historically high N deposition, summarizing >20 years of soil C data both during and post-termination of high N conditions. These findings will be critical in informing Earth System Models of how to parametrize projected forest C stocks in response to reduced anthropogenic N deposition.

Following the termination of the experimental N deposition treatment, C in the organic horizon (i.e., forest floor; Oe/Oa horizons) that accumulated over the course of this experiment (+18% change from ambient; Table 1) was no longer present within 5 years of treatment termination (Fig. 3), partially supporting our first hypothesis (H1a; condition 2; Fig. 1). The previously observed stability of the organic horizon C under the experimental N deposition treatment, and subsequent instability in ecosystem recovery that we present here, is ultimately defined by the balance between inputs (via litterfall and root biomass) and outputs (via decomposition) in the system. In our study system, over the 24-year experimental N deposition period, experimental N deposition did not significantly increase above-(litterfall) or below-ground (root biomass) inputs^4,11,27^ and similar findings have been observed at other long-term N-addition studies across the Eastern U.S.^5,10,28^. Accumulation of soil C across forests exposed to experimentally increased N deposition are thus attributed to the suppression of organic matter decomposition by soil microorganisms, not increased litter inputs^3–5,11,12,16,29^. In this system, the molecular mechanism mediating reduced organic matter decomposition was the transcriptional downregulation of genes encoding class II peroxidase enzymes, which completely oxidize complex polyphenols in plant detritus and soil organic matter to CO_2_^3^. Under lab conditions high levels of inorganic N, can suppress class II peroxidase genes in some Agaricomycetes, a response that can emerge within less than a week of high N exposure^30–33^. This transcriptional response may again underlie the observed ecosystem-level response, as class II peroxidase gene expression may increase after the alleviation of elevated N deposition to similar levels of expression observed under ambient conditions. Indeed, 5 years post-recovery, the activity potential of peroxidase enzymes in the former N deposition treatment increased (+22%, *P* = 0.03) as compared to the ambient condition (Fig. 5). Thus, it is plausible that the observed loss of organic horizon C 5-years post-termination of the experimental N deposition treatment is most likely the result of increases in organic matter decomposition by soil fungi rather than novel changes in inputs. While we do not have evidence that this transcriptional mechanism was responsible for carbon degradation in this study, this pattern is compelling and further work exploring the transcriptional responses of fungal class II peroxidases is warranted to assess this mechanism of C loss during forest recovery from historically high N deposition.

Estimations of microbial activity (i.e., microbial respiration and extracellular enzyme activity potential) support our claim that the rapid depletion of C in the organic horizon is the result of increases in decomposition. Following the termination of the N deposition treatment, we observed relative increases in microbial activity that partially support our hypothesis that organic horizon microbial activity would return to ambient conditions (H2a; condition 2; Fig. 1). During the experimental N deposition treatment, organic horizon microbial respiration was suppressed (−15%; Supplementary Table 1) in the N deposition treatment compared to ambient conditions^27^. Yet, 5-years post-termination of the N deposition treatment, there is no difference in microbial respiration in the organic horizon in the N deposition treatment and ambient conditions, demonstrating a relative increase in microbial respiration in the N deposition treatment following its termination. These observations align with models linking C and N dynamics that predict that when N is limiting in a system, increased inorganic N availability reduce respiration, whereas greater organic C availability increase it^34^. The primary mechanism underlying this dynamic has been attributed to microbial extracellular enzyme activity and the metabolic implications of that activity^15,35–38^. In recovery from historically high N deposition, microbial respiration likely increased due to the metabolic expense of extracellular enzyme production by soil microbes in order to break down the C to access adequate nutrients to meet their metabolic demands. We also observed increased lignin-modifying enzyme activity, as peroxidase activity potential increased in the N deposition treatment relative to ambient conditions (+22%, *P* = 0.03; Fig. 5). Taken together, our results suggest that 5 years into ecosystem recovery from historically high N deposition, the decay of more biochemically recalcitrant substrates that accumulated under experimental N deposition occurs via the increased production of extracellular enzymes, ultimately resulting in the depletion of the previously accumulated organic horizon C we have documented here.

Increased peroxidase enzyme activity in the former N deposition treatment may have altered interactions between lignolytic fungi and other organisms mediating organic matter decay. For example, during the 24-year period of the experimental N deposition treatment application, the downregulation of class II peroxidase genes occurred in the absence of a major change in community composition of fungi actively mediating lignin decay^39^. Instead, increases in the abundance of bacterial genes that produce enzymes which weakly oxidize polyphenols, such as those in lignin, were observed. This suggests that high N deposition elicited a community metabolic change from fungal decay of these substrates to their partial bacterial metabolism^40,41^. If this previously documented microbial response has reversed during ecosystem recovery, it could result in increased litter decay. Further work is necessary to determine whether this microbial mechanism contributes to the observed rapid loss of organic horizon C during ecosystem recovery from elevated N.

While three of the four sites we studied exhibited reduced organic horizon C following the termination of the N deposition treatment, the northernmost site A did not (Fig. 3; Extended Data Fig. 2). Interestingly, the persistence of the organic horizon C at site A is not explained by estimations of microbial activity. If the parameters of microbial activity we measured here do not regulate the amount of organic horizon C present in the system at site A an interesting question arises: what is regulating C cycling dynamics at northernmost site A? Long-term N addition studies find significant retention (>70%) of the added N in temperate forests^42,43^, with most of that retention occurring in the organic horizon^23,43,44^. During ecosystem recovery, soil inorganic N pools may recover to levels similar to that of ambient conditions within a few years^45^; whereas, total soil N may remain elevated for many years subsequently affecting other soil properties such as pH and nutrient concentrations^23^. As such, a potential explanation for the retention of the organic horizon C at site A may be the concurrent retention of N in the organic horizon. Though not statistically significant, we observed a +11% increase in organic horizon N in the N deposition treatment at site A (*P* = 0.17; Extended Data Fig. 5) as well as a sustained lower C:N ratio as compared to ambient conditions (Cohen’s d = -1.03 [-2.73, 0.76]; Extended Data Fig. 6), despite the N deposition treatment not receiving N-additions for 5-years. While previous work determined no differences in the extent of SOM decay across sites^46^, future work would benefit from a more high-resolution characterization of the SOM chemistry to elucidate the recovery trajectory and mechanisms underlying that of site A vs. the other sites.

High N retention in temperate forests may be attributable to climatic factors such as lower rates of precipitation that allow for longer contact time for N to be immobilized in the organic horizon^47^. Site A is the northernmost site of our study and experiences relatively low levels of precipitation and the lowest mean annual temperature (Fig. 2; Supplemental Table 2). Therefore, we suspect differences in climate at site A compared to the three southernmost sites may indirectly impact decomposition rates, suggesting recovery from historically high N deposition may be governed by differences in climatic factors, though we do not have the evidence to directly support this at this time. Climate is one of the primary controls over soil organic C dynamics^48–52^ by affecting microbial community composition and activity that regulate C cycling in forests globally^53^. As climate change and recovery from historically high N deposition progress concomitantly, it will be increasingly imperative to disentangle the effects of changing N availability and changing climate on forest C cycling and storage.

Despite the rapid and dynamic response of the organic horizon C to recovery from the N deposition treatment, the mineral soil C pool is recovering at a slower rate (Fig. 4a), partially supporting H1b (condition 1; Fig. 1). This finding aligns with a hysteretic model of recovery whereby the state of a property (e.g., soil C) lags behind changes in the effect causing it (e.g., reduced N availability)^23^. In ecological systems, this occurs when the output is not a strict function of the corresponding input due to the complex suite of variables that affect biological activity^23,54^. In this study, mineral soil C 5-years into recovery from high N remains elevated in the former N deposition treatment compared to ambient conditions (+11% change from ambient), but the increase is no longer statistically significant (*P* = 0.31; Fig. 4b). This is supported by our finding that peroxidase activity potential remains suppressed, though not statistically significant, in the mineral soil (−18% change from ambient, *P* = 0.06; Extended Data Fig. 4), partially supporting hypothesis H2b (condition 1; Fig. 1) and providing a potential mechanism explaining the marginal persistence of C. However, the soils in our experiment are sandy (∼85%) spodosols and thus do not generally have a high capacity to retain mineral soil C and are more likely than finer textured soils to experience faster changes in C concentrations in response to disturbance^55,56^. Therefore, we suspect that the soil physical properties of our system control the status of mineral soil C in recovery. Ultimately, the data we have obtained 5-years post-termination of the N deposition treatment exhibit marginal responses from the mineral soil C pool, though we speculate the degree of response will change over time. If the mineral soil peroxidase enzyme activity of the N deposition treatment nears that of ambient conditions like we observed in the forest floor, we predict mineral soil C losses will similarly occur.

## CONCLUSIONS

Our results demonstrate that 5-years into recovery from experimental N deposition, the organic horizon C that accumulated has been depleted in the southernmost three sites due to increases in microbial activity, whereas the accumulated organic horizon C in the northernmost site has persisted. As recovery in our study system progresses, we hypothesize the following three scenarios will occur: the first is that we expect organic horizon C levels in the former N deposition treatment at the northernmost site to become more similar to organic horizon C levels under ambient conditions, possibly eventually exhibiting reductions similar to the three southernmost sites. The second, is that given the loss of mineral soil C we observed, we expect the mineral soil pool to continue to lose the C that accumulated under experimental N deposition (Fig. 6). Finally, we do not expect the three southernmost sites to continue to lose organic horizon C. Rather, we suspect the system is in a transitory period and exhibiting a perturbation following disturbance that could manifest as early warning signs for destabilization as this globally relevant terrestrial C sink moves towards a new equilibrium point (Fig. 6)^52^. We suspect the organic horizon dynamics we have presented may occur in other temperate forest systems and long-term N addition studies, though the degree of change will likely be different depending on the dominant tree species present, soil physical properties, and climatic factors. Additional monitoring of this and other long-term experimental N studies in recovery will be necessary to accurately understand forest C response, which will be crucial in modeling future forest C stocks under environmental change. If other terrestrial ecosystems respond similarly to reductions in atmospheric N deposition, the global terrestrial C sink in the northern hemisphere will decline and subsequently enhance climate warming.

**Figure 6.**
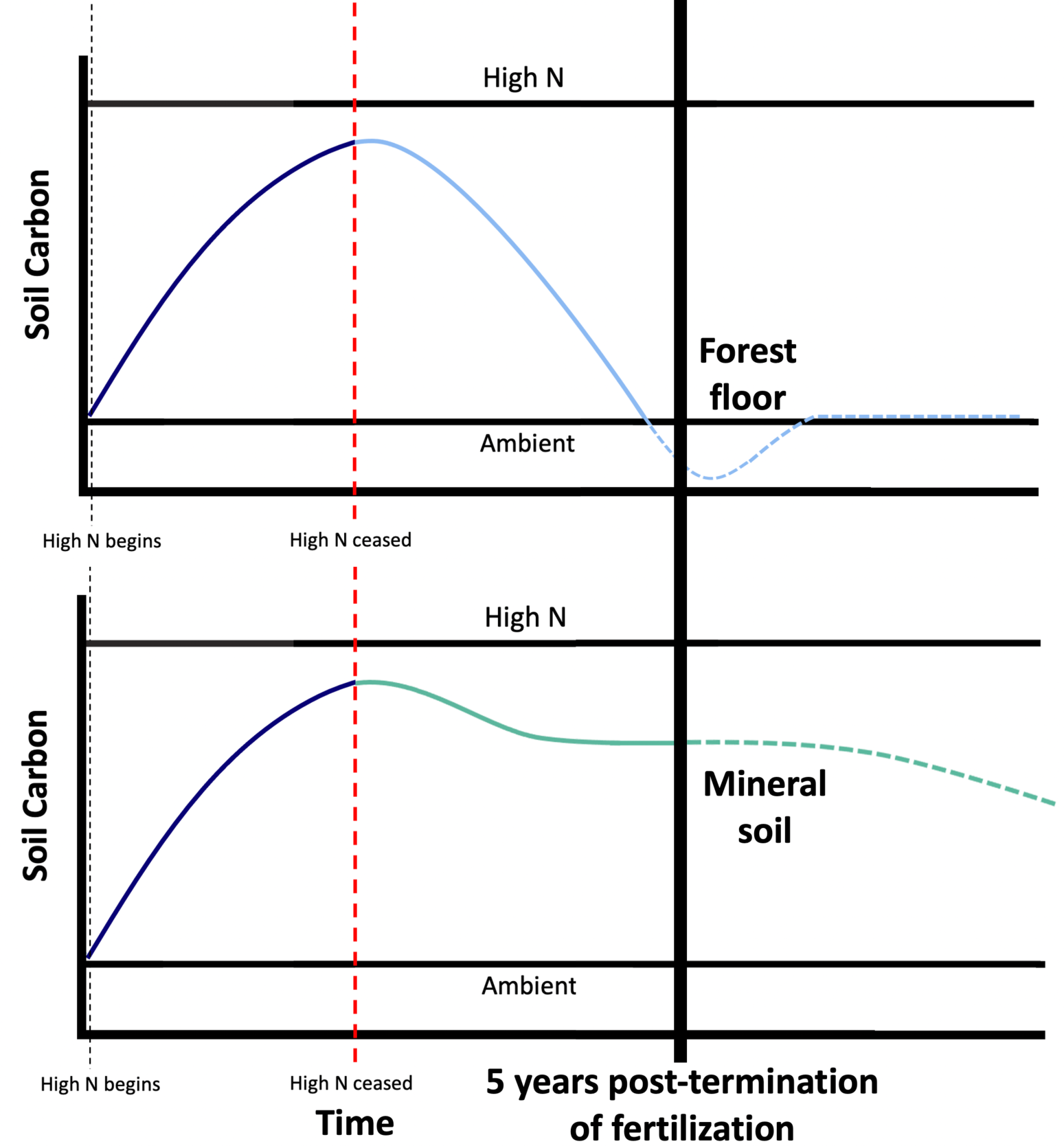
Proposed future recovery trajectory of organic horizon (top) and mineral soil (bottom) C from historically high N deposition. At 5-years post-termination of the N deposition treatment (as indicated by the solid black line), soil C that accumulated in the organic horizon (i.e., forest floor) exhibits not only a return to the ambient condition state (condition 2; Fig. 1), but an additional deficit relative to ambient conditions. We speculate that organic horizon C will eventually return to that of the ambient state. Mineral soil C that accumulated during high N deposition is also being lost but at a slower rate. We speculate that mineral soil C will continue to be lost, potentially reaching a new steady state (condition 3; see Fig. 1).

## METHODS

### Site description and experimental design

The Michigan Nitrogen Deposition Gradient Study was established in 1987 to examine the effects of climate and atmospheric N deposition on ecosystem processes. Spanning a 500-km distance, the study area in Michigan, USA, consists of four edaphically and floristically matched sugar maple (*Acer saccharum*) dominated northern hardwood forest stands on sandy spodosol soil (Typic Haplorthods of the Kalkaska series) (Fig. 2; Supplementary Table 2). The stands span a natural climatic gradient that encompasses the full latitudinal range of northern hardwood forests in eastern North America and the Upper Great Lakes Regio^57^, allowing us to generalize our findings across this widespread and important ecosystem in North America (Supplementary Table 2). At each site (n=4), there are six 30-m by 30-m plots, three of which received experimental N additions in the form of 30 kg NO_3_^—^N ha^-1^ yr^-1^ beginning in 1994 and three of which receive ambient atmospheric N deposition. The experimental N addition treatment was applied annually over a 24-year period and was terminated following the final application in the summer of 2017. All data obtained prior to 2022 were generated using similar sampling techniques and analytical methods to those described here and can be found in detail in Pregitzer et al. (Ref. 11).

### Field sampling and sample processing

We collected samples in September of 2022. At each site, 15 samples of organic horizon (i.e., forest floor) and 10 samples of mineral soil were collected at random in each plot (n=6, per site) and composited in the field to produce 2 homogenized forest floor samples and 1 homogenized mineral soil sample per plot. Freshly fallen litter (Oi horizon) was removed by hand and the underlying Oe and Oa horizons were collected using a 10-cm by 10-cm PVC square and cut with a serrated knife to the mineral soil surface. Of the 15 forest floor samples collected, 10 were composited to yield one homogenized sample that would later be air dried and the remaining 5 samples were composited to yield one homogenized sample that would be used for metrics of biological activity. At 10 of the 15 locations where the organic horizon was sampled, mineral soil samples were taken from the center of the PVC square using a 2-cm-diameter soil corer (10-cm-depth; A horizon) and were homogenized by hand in the field to produce one mineral soil sample per plot. Samples were then immediately put on ice for transport to the University of Wisconsin-Madison. Upon return to the lab, forest floor samples were sifted by hand to remove materials not classified as Oe/Oa horizon. Mineral soil samples were passed through a 2-mm sieve to remove rocks and roots. During and after this initial processing, all samples were stored at 4*°*C in the lab.

### Soil C and N content

To assess if the SOM that had accumulated in the organic horizon and mineral soil in the experimental N deposition treatment persists following the termination of N fertilization, forest floor mass and organic horizon and mineral soil C and N content were measured. Forest floor samples were air dried and weighed daily until their mass equilibrated at which time final measurements of forest floor mass were obtained. Total C and N of forest floor and mineral soil samples was measured by flash combustion on a FlashEA 1112 Elemental Analyzer (Thermo Scientific, Waltham, MA, USA). Individual sample bulk density was used to calculate volumetric soil C stocks (g C m^-2^).

### Microbial Respiration

To measure microbial respiration of organic horizon and mineral soil samples, a laboratory incubation experiment was performed using methods derived from Franzluebbers et al. (Refs. 58 & 59) and Wade et al. (Ref. 60). Twenty grams of air-dried mineral soil or two grams of air-dried forest floor were placed in 0.946-L air-tight containers. While working in the fume hood to ensure uniform initial levels of CO_2_ in each sample container, soil moisture content was brought to 50% soil water holding capacity (WHC) by dispensing Milli-Q water in a circular motion to prevent splashing and minimize disturbance effects on biological activity^61^. WHC was calculated as grams of water held per gram of soil as described in Wade et al. (Ref. 60). After bringing the moisture of each sample to 50% WHC, containers were sealed with air-tight lids containing a rubber septum and pre-incubated for 7-days, during which they were stored in a dark room with air temperature kept at 20°C. Following the 7-day pre-incubation period, each sample container was uncapped for 10-minutes in the fume hood to allow container headspace to equilibrate with ambient atmospheric CO_2_ levels. Additional Milli-Q water was added when needed to ensure soil moisture remained at 50% WHC. Sample containers were sealed and stored again, and headspace gas was sampled after 24, 48, and 72 hours. Headspace CO_2_ samples were drawn by hand by taking a 10 ml sample using a 30 ml polypropylene syringe fitted with a 21-gauge needle. Samples were then injected over a 30-second period per sample into an EGM-5 Infrared Gas Analyzer (IRGA) (PP Systems, Amesbury, MA, USA). Values obtained from the 72-hour sampling period were used to calculate C respired as g C g soil^-1^ s^-1^.

### Extracellular Enzyme Activity

We measured extracellular enzymes with key roles in mediating SOM decomposition and soil C storage using methods from Saiya-Cork et al. (Ref. 62), which have consistently been used to assess changes in enzyme activity in this long-term experimental study. The enzymes, β-glucosidase (BG), polyphenol oxidase (PPO), and peroxidase (PER), catalyze the metabolism of two common plant litter compounds, hemicellulose (BG) and lignin (PPO and PER), are highly associated with soil C^15^, and have consistently exhibited significant changes to experimental N deposition in our study system^3,36,37,63,64^. Two grams of field-fresh mineral soil or one gram of field-fresh forest floor were combined with 125 ml (for mineral soil) or 150 ml (for forest floor) of sodium acetate buffer (pH 5) and homogenized for 1 minute using a BioHomogenizer (BioSpec Products, Bartlesville, OK, USA) to produce a homogenized soil slurry. All soil slurries were continuously stirred to ensure homogeneity and plated on 96-well plates within 30-minutes. BG activity potential was measured following a 5-hour incubation period at 20°C using a fluorometric assay with 355 nm excitation and 460 nm emission filters with 4-Methylumbelliferyl β-D-glucopyranoside as the substrate. PPO and PER activity potential were measured following an 18-hour incubation period at 20°C using a colorimetric assay by measuring absorbance at 460 nm with L-dihydroxyphenylalanine (L-DOPA) as the substrate and the addition of hydrogen peroxide to PER assays. All enzyme assays were performed on a FLUOstar Omega microplate reader (BMG Labtech, Ortenberg, Germany). Enzyme activity potential was expressed in units of nmol substrate oxidized h^-1^ g^-1^.

### Statistical analyses

We calculated standardized effect sizes using the effectsize package (v0.8.5.)^65^ in R by calculating Cohen’s d with a 95% confidence interval to compare the effect of experimental N treatment, and its termination, on soil C content in the organic and mineral soil horizons at each site across time since the beginning of our long-term experiment (1994-2022)^66^. To assess recovery at a finer resolution, we used two-way analysis of variance (ANOVA) with a site by treatment interaction to determine whether historic experimental N-additions, herein referred to as the N deposition treatment, had a significant effect on the SOM content, mass, and microbial respiration in organic horizon and mineral soil pools 5-years post-termination of experimental N additions. In this system, we previously found that soil water content has a significant influence on enzyme activity^37^, and as such, we used two-way analysis of co-variance (ANCOVA) with a site by treatment interaction to determine whether historic experimental N additions had a significant effect on extracellular enzyme activity potential 5-years post-termination of the N deposition treatment while controlling for soil moisture at the sample level. ANCOVA was performed using the rstatix package (v0.7.2)^67^ in R. When applicable, least square means was used to estimate the marginal means of extracellular enzyme activity potential adjusted for soil moisture and among-group means were compared using Tukey’s HSD. ANCOVAs were performed on the estimated marginal means, but raw values are presented. All data processing, statistical analyses, and visualizations were performed and produced in R (v4.2.2)^68^ and RStudio (version 2022.12.0+353)^69^.

## ACKNOWLEDGEMENTS

We thank Jeremy Fleck, Emily Sautebin, Alessia Fucentese, Annalise Keaton, Caleb Spector, and Annalisa Stevenson for their help in the lab and field. We also wish to thank Thea Whitman (and lab) and Will Argiroff for their thoughtful feedback on this work. This material is based upon work supported by the National Science Foundation Graduate Research Fellowship Program under Grant No. DGE-2137424 and by USDA National Institute of Food and Agriculture, Hatch project accession numbers 1029318.

---

**Extended Data Figure 1.**
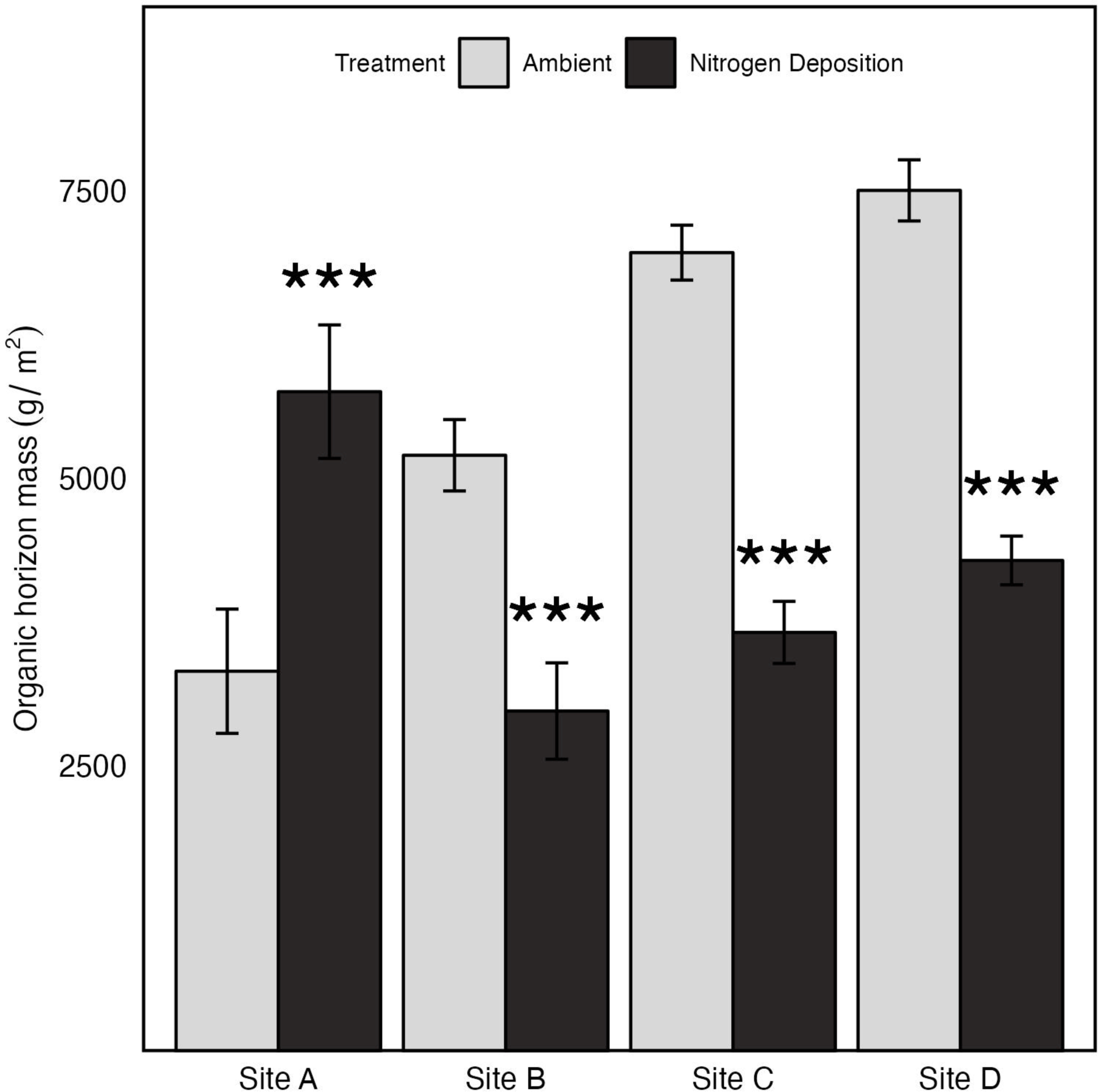
The effect of the historically high N deposition and site on the mass of the organic horizon from samples collected in September 2022, 5-years post-termination of the N deposition treatment. Site values represent treatment means ± standard error (n=3). *** *P* < 0 .001.

**Extended Data Figure 2.**
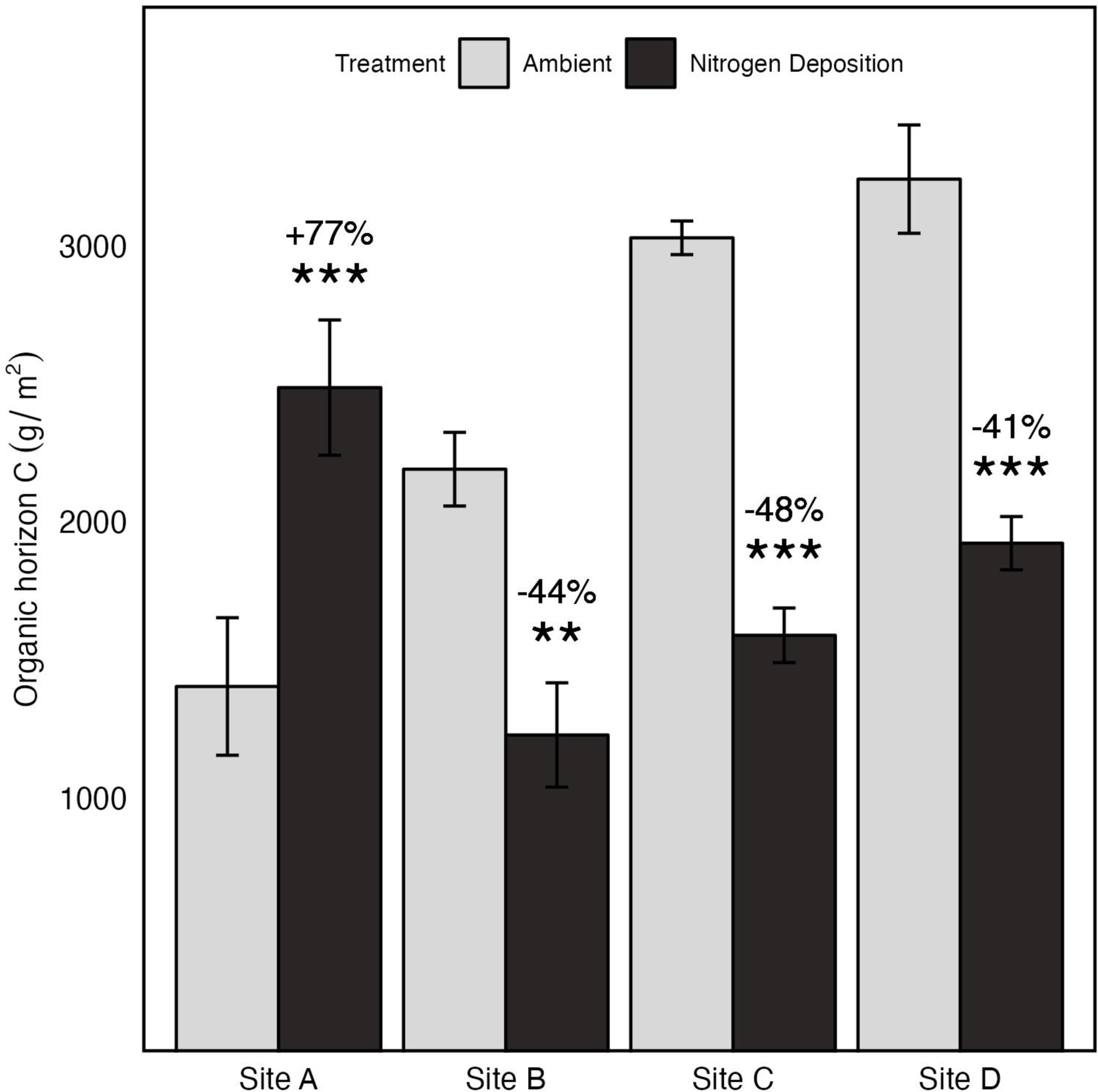
The effect of the historically high N deposition and site on organic horizon C from samples collected in September 2022, 5-years post-termination of the N deposition treatment. Site values represent treatment means ± standard error (n=3). *** *P* < 0 .001 and ** *P* < 0.01. Organic horizon C values in the N deposition treatment percent change from ambient are reported.

**Extended Data Figure 3.**
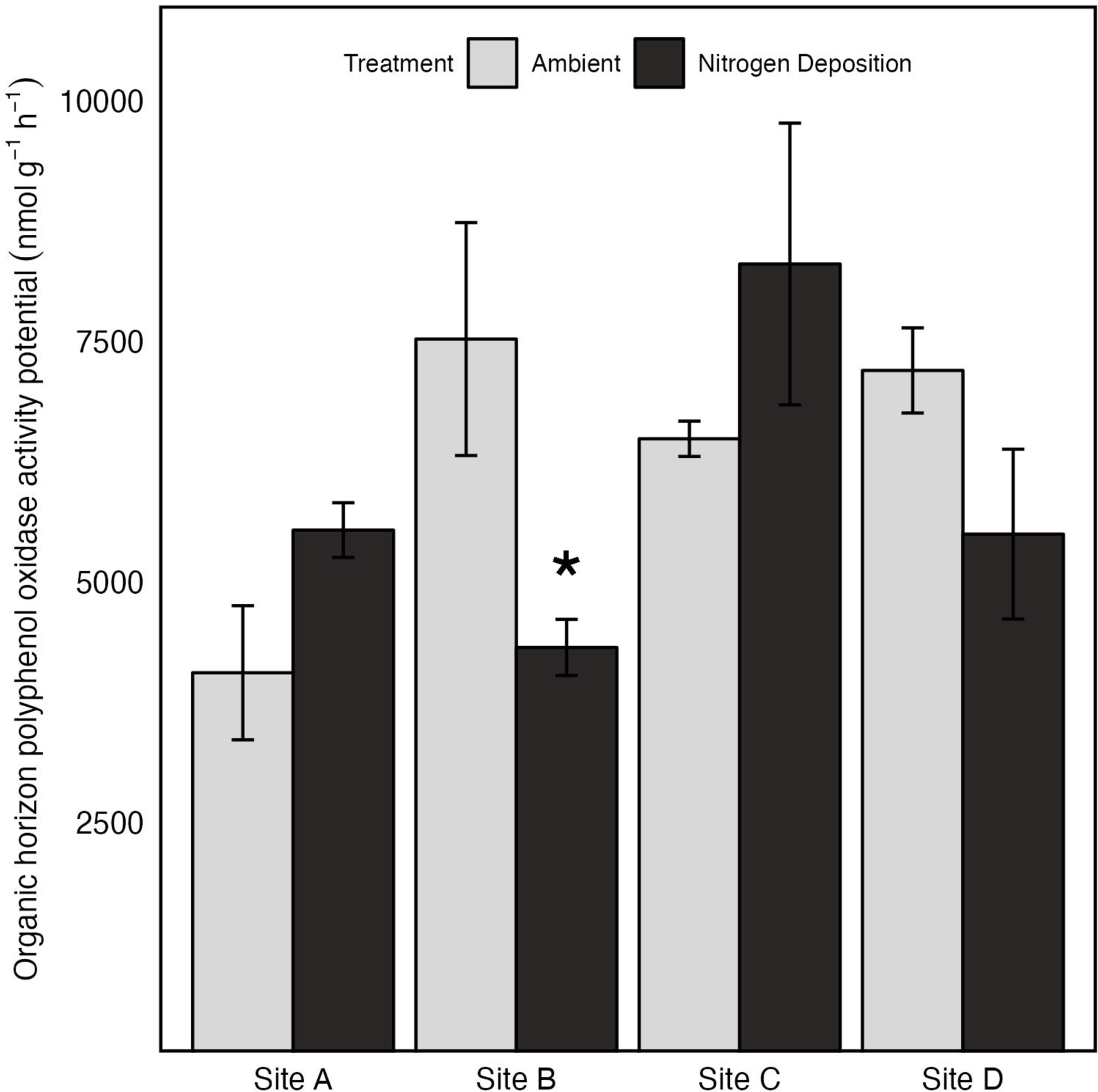
The effect of historically high N deposition on organic horizon polyphenol oxidase (PPO) activity values from samples collected in September 2022, 5-years post-termination of the N deposition treatment. ANOVA was performed on estimated marginal means, but raw values are presented. Site values represent treatment means ± standard error (n=3). * *P* < 0 .05.

**Extended Data Figure 4.**
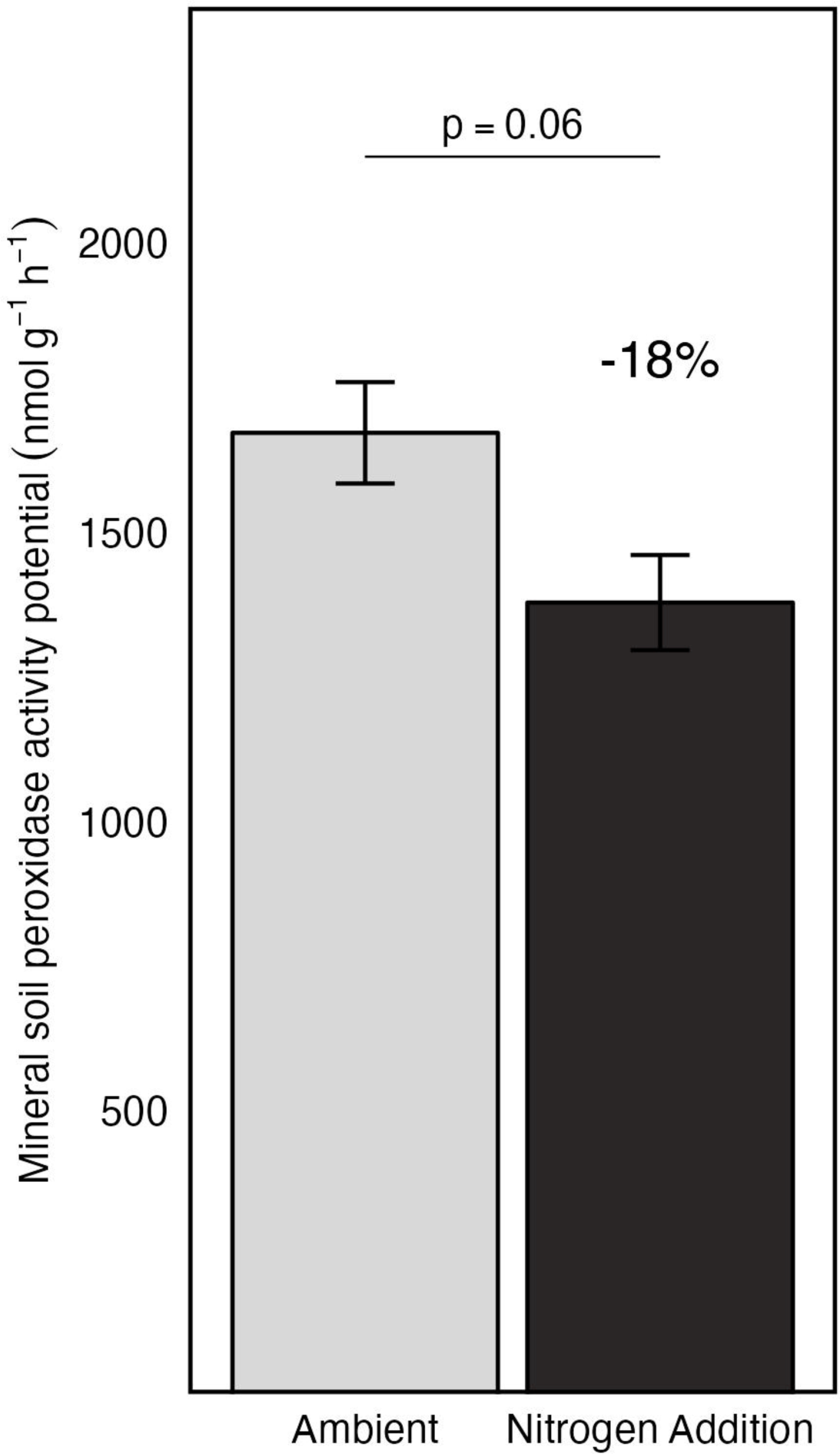
The effect of historically high N deposition on mineral soil peroxidase (PER) activity values from samples collected in September 2022, 5-years post-termination of the N deposition treatment. ANOVA was performed on estimated marginal means, but raw values are presented. Values represent treatment means ± standard error (n=12). Site × Treatment *P* > 0.05. Mineral soil PER values in the N deposition treatment percent change from ambient is reported.

**Extended Data Figure 5.**
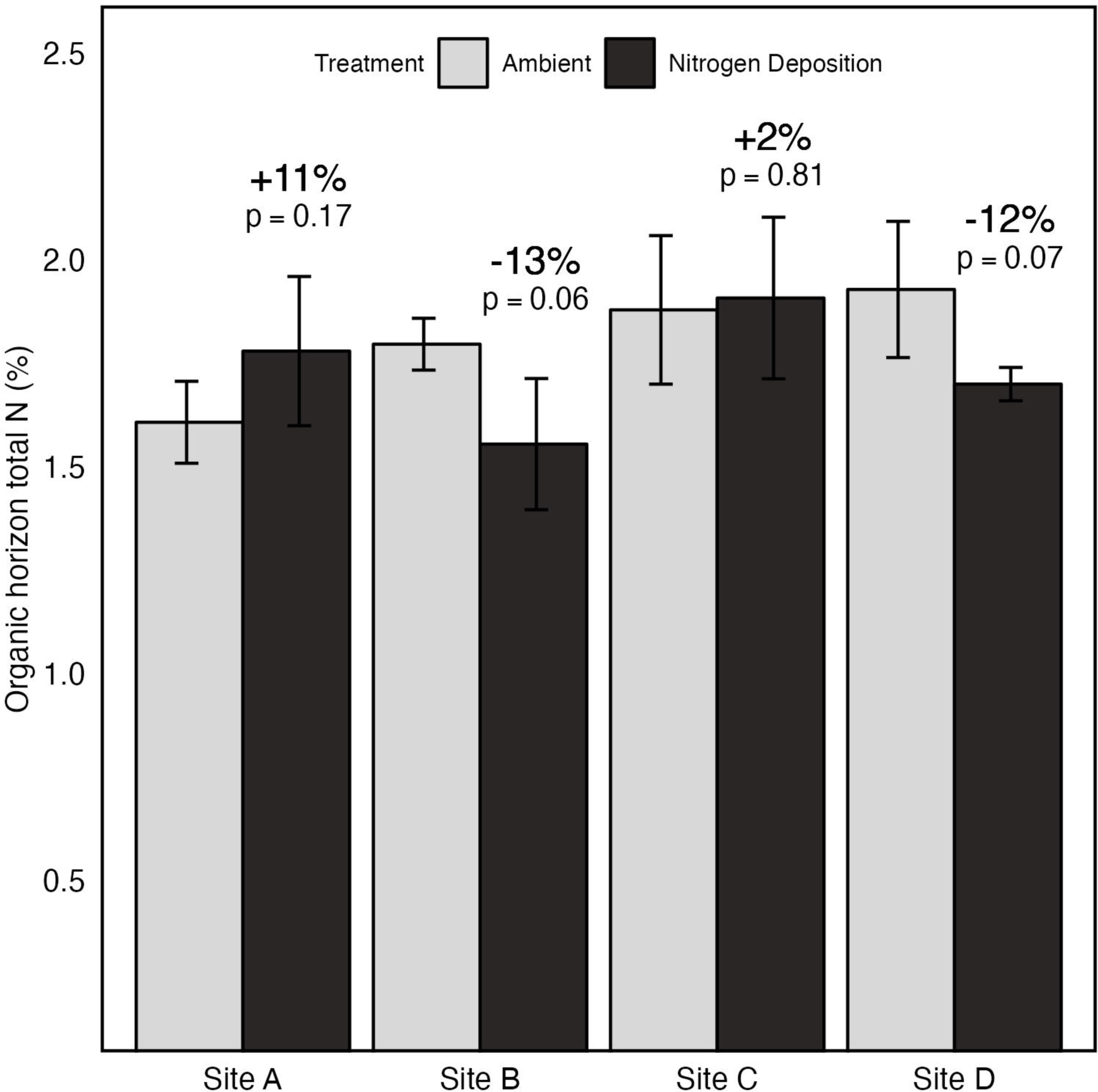
The effect of the historically high N deposition and site on organic horizon N from samples collected in September 2022, 5-years post-termination of the N deposition treatment. Site values represent treatment means ± standard error (n=3). Organic horizon N values in the N deposition treatment percent change from ambient are reported.

**Extended Data Figure 6.**
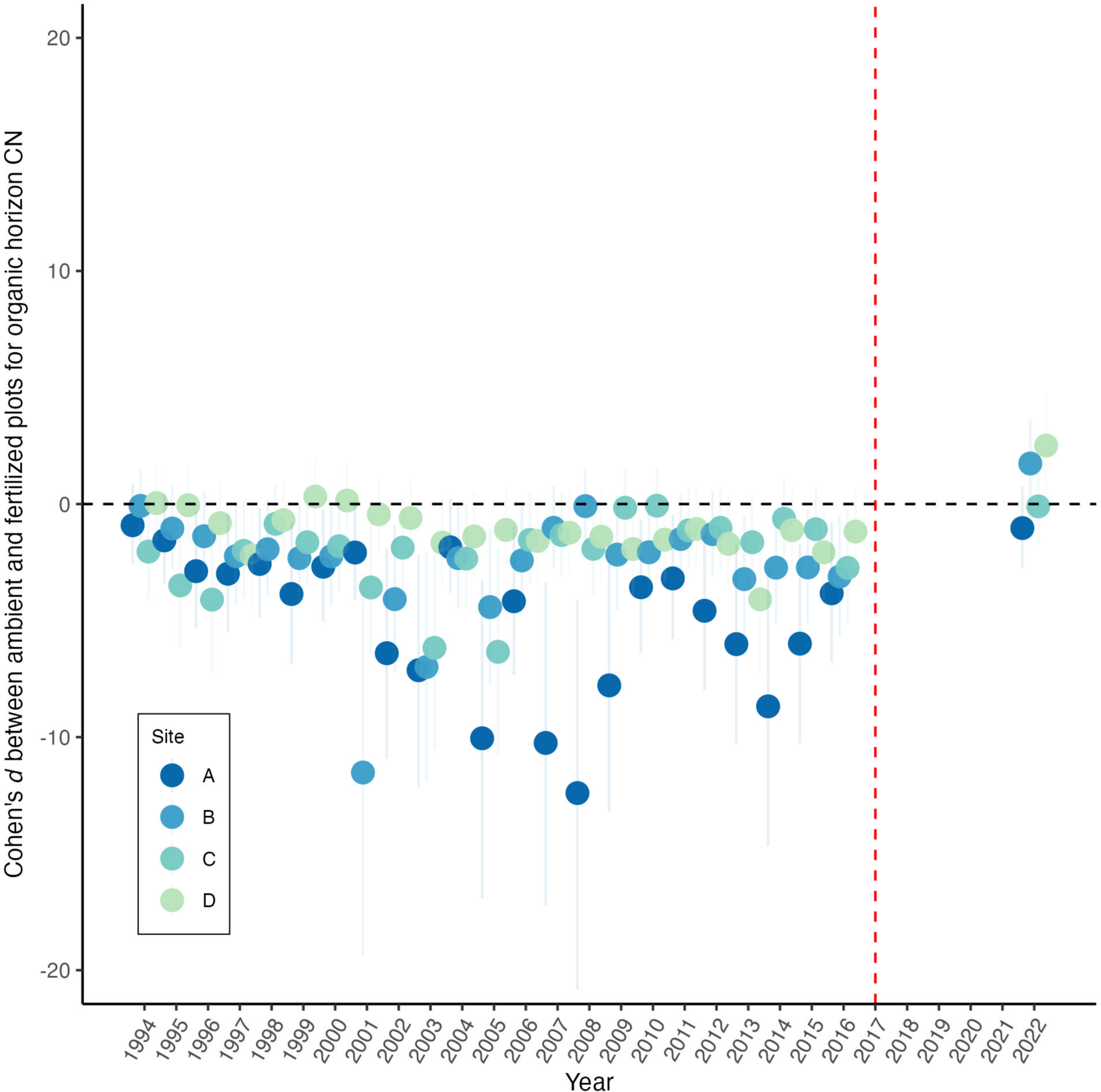
Standardized effect sizes (Cohen’s d) with 95% confidence intervals of the N deposition treatment on organic horizon C:N throughout the long-term experiment. The N deposition treatment was applied annually from 1994-2017. The red vertical line indicates the first year in which no treatment was applied. Points are colored by site (n = 4). Points below zero indicate a lower organic horizon C:N ratio in the N deposition treatment compared to the ambient treatment. Points above zero indicate a higher C:N ratio in the organic horizon in the N deposition treatment compared to the ambient treatment.

**Supplementary Table 1.**
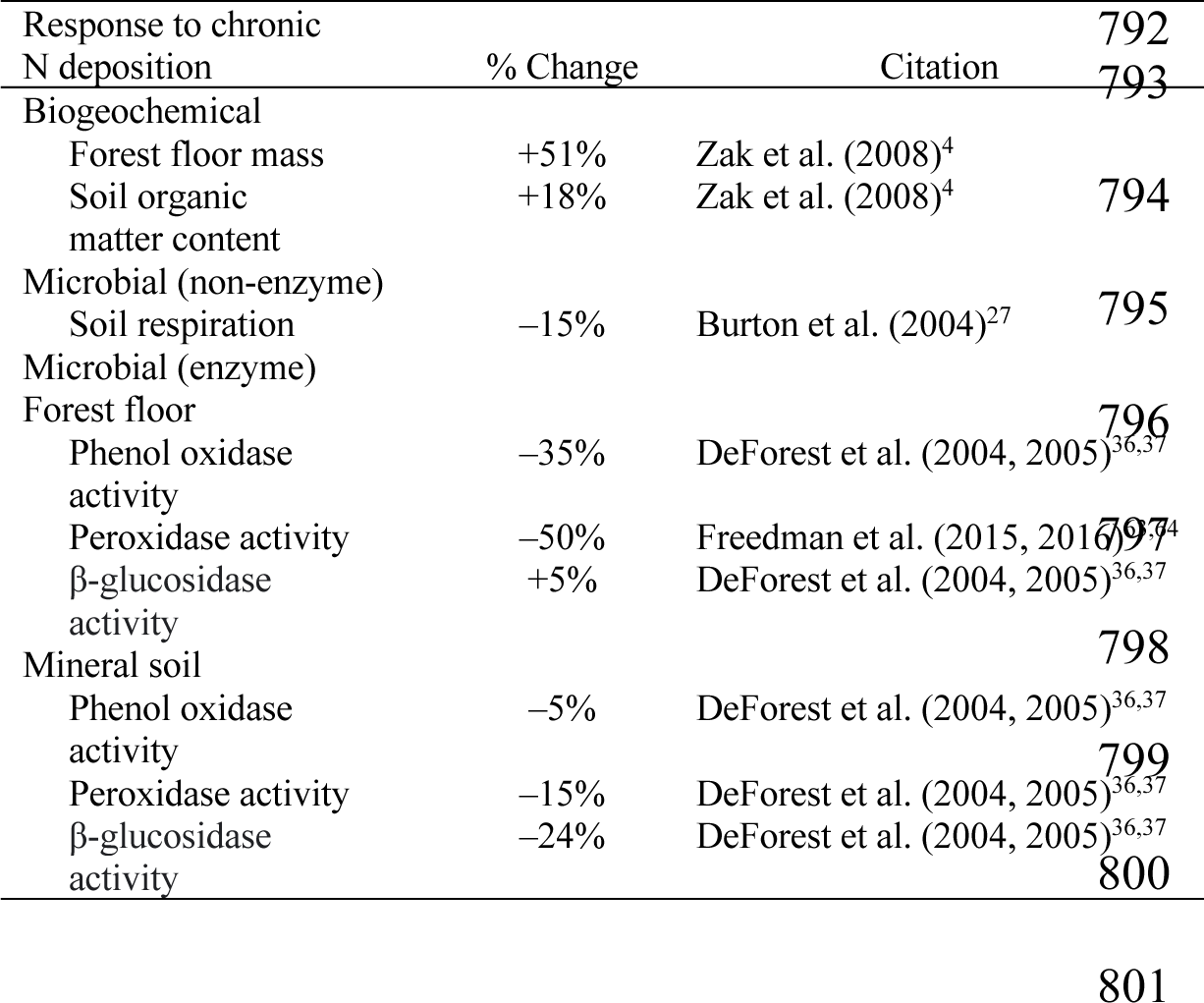
Relative changes from ambient exhibited in response to experimental N deposition resulting in increased C storage and reduced cycling of C in forest floor and surface mineral soil. Non-enzymatic responses are summarized for forest floor (Oe/Oa), where mineral soil responses have exhibited similar trends. With the exception of forest floor β-glucosidase and mineral soil phenol oxidase and peroxidase activities, all positive and negative responses are statistically significant (*P* < 0.05).

**Supplementary Table 2.** Site, climate, vegetation, and soil characteristics of the four northern hardwood forest stands in Michigan, USA (see Fig. 2). The sites exist across a natural north-south climatic gradient, yet are similar in age, plant composition, and soil development.

| Characteristic | Site A | Site B | Site C | Site D |
| --- | --- | --- | --- | --- |
| *Location* |  |  |  |  |
| Latitude (N) | 46°52' | 45°33' | 44°23' | 43°40' |
| Longitude (W) | 88°53' | 84°52' | 85°50' | 86°09' |
| *Climate* |  |  |  |  |
| Mean annual temperature (°C) | 4.7 | 6.0 | 6.9 | 7.6 |
| Mean annual precipitation (mm) | 873 | 871 | 888 | 812 |
| Ambient wet + dry total N deposition (g m<sup>-2</sup> yr<sup>-1</sup>) | 0.68 | 0.91 | 1.17 | 1.18 |
| Growing season length (days) | 134 | 150 | 154 | 157 |
| *Vegetation* |  |  |  |  |
| Overstory biomass (Mg ha<sup>-1</sup>) | 261 | 261 | 274 | 234 |
| *Acer saccharum* biomass (Mg ha<sup>-1</sup>) | 237 | 224 | 216 | 201 |
| *Soil* |  |  |  |  |
| Soil texture (0-10cm) (%sand-%silt-%clay) | 75:22:3 | 89:9:2 | 89:9:2 | 87:10:3 |

## REFERNCES

1. Keenan, T. F., Prentice, I. C., Canadell, J. G., Williams, C. A., Wang, H., Raupach, M., & Collatz, G. J. (2016). Recent pause in the growth rate of atmospheric CO2 due to enhanced terrestrial carbon uptake. Nature Communications, 7(1), 13428. 10.1038/ncomms13428

2. Magnani, F., Mencuccini, M., Borghetti, M., Berbigier, P., Berninger, F., Delzon, S., Grelle, A., Hari, P., Jarvis, P. G., Kolari, P., Kowalski, A. S., Lankreijer, H., Law, B. E., Lindroth, A., Loustau, D., Manca, G., Moncrieff, J. B., Rayment, M., Tedeschi, V., … Grace, J. (2007). The human footprint in the carbon cycle of temperate and boreal forests. Nature, 447(7146), 849–851. 10.1038/nature05847

3. Zak, D. R., Argiroff, W. A., Freedman, Z. B., Upchurch, R. A., Entwistle, E. M., & Romanowicz, K. J. (2019). Anthropogenic N deposition, fungal gene expression, and an increasing soil carbon sink in the Northern Hemisphere. Ecology, 100(10). 10.1002/ecy.2804

4. Zak, D. R., Holmes, W. E., Burton, A. J., Pregitzer, K. S., & Talhelm, A. F. (2008). SIMULATED ATMOSPHERIC NO 3 − DEPOSITION INCREASES SOIL ORGANIC MATTER BY SLOWING DECOMPOSITION. Ecological Applications, 18(8), 2016–2027. 10.1890/07-1743.1

5. Frey, S. D., Ollinger, S., Nadelhoffer, K., Bowden, R., Brzostek, E., Burton, A., Caldwell, B. A., Crow, S., Goodale, C. L., Grandy, A. S., Finzi, A., Kramer, M. G., Lajtha, K., LeMoine, J., Martin, M., McDowell, W. H., Minocha, R., Sadowsky, J. J., Templer, P. H., & Wickings, K. (2014). Chronic nitrogen additions suppress decomposition and sequester soil carbon in temperate forests. Biogeochemistry, 121(2), 305–316. 10.1007/s10533-014-0004-0

6. Huang, X., Terrer, C., Dijkstra, F. A., Hungate, B. A., Zhang, W., & van Groenigen, K. J. (2020). New soil carbon sequestration with nitrogen enrichment: A meta-analysis. Plant and Soil, 454(1–2), 299–310. 10.1007/s11104-020-04617-x

7. Knorr, M., Frey, S. D., & Curtis, P. S. (2005). NITROGEN ADDITIONS AND LITTER DECOMPOSITION: A META-ANALYSIS. Ecology, 86(12), 3252–3257. 10.1890/05-0150

8. Liu, L., & Greaver, T. L. (2010). A global perspective on belowground carbon dynamics under nitrogen enrichment: Belowground C dynamics under N enrichment. Ecology Letters, 13(7), 819–828. 10.1111/j.1461-0248.2010.01482.x

9. Lovett, G. M., Arthur, M. A., Weathers, K. C., Fitzhugh, R. D., & Templer, P. H. (2013). Nitrogen Addition Increases Carbon Storage in Soils, But Not in Trees, in an Eastern U.S. Deciduous Forest. Ecosystems, 16(6), 980–1001. 10.1007/s10021-013-9662-3

10. Magill, A. H., & Aber, J. D. (1998). Long-term effects of experimental nitrogen additions on foliar litter decay and humus formation in forest ecosystems. Plant and Soil, 203(2), 301–311. 10.1023/A:1004367000041

11. Pregitzer, K. S., Burton, A. J., Zak, D. R., & Talhelm, A. F. (2008). Simulated chronic nitrogen deposition increases carbon storage in Northern Temperate forests: NITROGEN DEPOSITION INCREASES CARBON STORAGE. Global Change Biology, 14(1), 142–153. 10.1111/j.1365-2486.2007.01465.x

12. Bowden, R. D., Wurzbacher, S. J., Washko, S. E., Wind, L., Rice, A. M., Coble, A. E., Baldauf, N., Johnson, B., Wang, J., Simpson, M., & Lajtha, K. (2019). Long-term Nitrogen Addition Decreases Organic Matter Decomposition and Increases Forest Soil Carbon. Soil Science Society of America Journal, 83(S1). 10.2136/sssaj2018.08.0293

13. Myneni, R. B., Dong, J., Tucker, C. J., Kaufmann, R. K., Kauppi, P. E., Liski, J., Zhou, L., Alexeyev, V., & Hughes, M. K. (2001). A large carbon sink in the woody biomass of Northern forests. Proceedings of the National Academy of Sciences, 98(26), 14784–14789. 10.1073/pnas.261555198

14. Schimel, D., Stephens, B. B., & Fisher, J. B. (2015). Effect of increasing CO 2 on the terrestrial carbon cycle. Proceedings of the National Academy of Sciences, 112(2), 436–441. 10.1073/pnas.1407302112

15. Chen, J., Luo, Y., van Groenigen, K. J., Hungate, B. A., Cao, J., Zhou, X., & Wang, R. (2018). A keystone microbial enzyme for nitrogen control of soil carbon storage. Science Advances, 4(8), eaaq1689. 10.1126/sciadv.aaq1689

16. Janssens, I. A., Dieleman, W., Luyssaert, S., Subke, J.-A., Reichstein, M., Ceulemans, R., Ciais, P., Dolman, A. J., Grace, J., Matteucci, G., Papale, D., Piao, S. L., Schulze, E.-D., Tang, J., & Law, B. E. (2010). Reduction of forest soil respiration in response to nitrogen deposition. Nature Geoscience, 3(5), 315–322. 10.1038/ngeo844

17. Lu, X., Vitousek, P. M., Mao, Q., Gilliam, F. S., Luo, Y., Turner, B. L., Zhou, G., & Mo, J. (2021). Nitrogen deposition accelerates soil carbon sequestration in tropical forests. Proceedings of the National Academy of Sciences, 118(16), e2020790118. 10.1073/pnas.2020790118

18. Erisman, J. W., Grennfelt, P., & Sutton, M. (2003). The European perspective on nitrogen emission and deposition. Environment International, 29(2–3), 311–325. 10.1016/S0160-4120(02)00162-9

19. Du, E. (2016). Rise and fall of nitrogen deposition in the United States. Proceedings of the National Academy of Sciences, 113(26). 10.1073/pnas.1607543113

20. Li, Y., Schichtel, B. A., Walker, J. T., Schwede, D. B., Chen, X., Lehmann, C. M. B., Puchalski, M. A., Gay, D. A., & Collett, J. L. (2016). Increasing importance of deposition of reduced nitrogen in the United States. Proceedings of the National Academy of Sciences, 113(21), 5874–5879. 10.1073/pnas.1525736113

21. Lloret, J., & Valiela, I. (2016). Unprecedented decrease in deposition of nitrogen oxides over North America: The relative effects of emission controls and prevailing air-mass trajectories. Biogeochemistry, 129(1–2), 165–180. 10.1007/s10533-016-0225-5

22. Mason, R. E., Craine, J. M., Lany, N. K., Jonard, M., Ollinger, S. V., Groffman, P. M., Fulweiler, R. W., Angerer, J., Read, Q. D., Reich, P. B., Templer, P. H., & Elmore, A. J. (2022). Evidence, causes, and consequences of declining nitrogen availability in terrestrial ecosystems. Science, 376(6590), eabh3767. 10.1126/science.abh3767

23. Gilliam, F. S., Burns, D. A., Driscoll, C. T., Frey, S. D., Lovett, G. M., & Watmough, S. A. (2019). Decreased atmospheric nitrogen deposition in eastern North America: Predicted responses of forest ecosystems. Environmental Pollution, 244, 560–574. 10.1016/j.envpol.2018.09.135

24. Gilliam, F. S. (2021). Response of Temperate Forest Ecosystems under Decreased Nitrogen Deposition: Research Challenges and Opportunities. Forests, 12(4), 509. 10.3390/f12040509

25. Carrara, J. E., Fernandez, I. J., & Brzostek, E. R. (2022). Mycorrhizal type determines root– microbial responses to nitrogen fertilization and recovery. Biogeochemistry, 157(2), 245–258. 10.1007/s10533-021-00871-y

26. Ollinger, S. V., Smith, M. L., Martin, M. E., Hallett, R. A., Goodale, C. L., & Aber, J. D. (2002). Regional Variation in Foliar Chemistry and N Cycling among Forests of Diverse History and Composition. Ecology, 83(2), 339. 10.2307/2680018

27. Burton, A. J., Pregitzer, K. S., Crawford, J. N., Zogg, G. P., & Zak, D. R. (2004). Simulated chronic NO 3 − deposition reduces soil respiration in northern hardwood forests: CHRONIC NO 3 − ADDITIONS REDUCE SOIL RESPIRATION. Global Change Biology, 10(7), 1080–1091. 10.1111/j.1365-2486.2004.00737.x

28. Eastman, B. A., Adams, M. B., Brzostek, E. R., Burnham, M. B., Carrara, J. E., Kelly, C., McNeil, B. E., Walter, C. A., & Peterjohn, W. T. (2021). Altered plant carbon partitioning enhanced forest ecosystem carbon storage after 25 years of nitrogen additions. New Phytologist, 230(4), 1435–1448. 10.1111/nph.17256

29. Adams, M. B., & Angradi, T. R. (1996). Decomposition and nutrient dynamics of hardwood leaf litter in the Fernow Whole-Watershed Acidification Experiment. Forest Ecology and Management, 83(1–2), 61–69. 10.1016/0378-1127(95)03695-4

30. Van Der Woude, M. W., Boominathan, K., & Reddy, C. A. (1993). Nitrogen regulation of lignin peroxidase and manganese-dependent peroxidase production is independent of carbon and manganese regulation in Phanerochaete chrysosporium. Archives of Microbiology, 160(1), 1–4. 10.1007/BF00258138

31. Tien, M., & Tu, C.-P. D. (1987). Cloning and sequencing of a cDNA for a ligninase from Phanerochaete chrysosporium. Nature, 326(6112), 520–523. 10.1038/326520a0

32. Brown, J. A., Alic, M., & Gold, M. H. (1991). Manganese peroxidase gene transcription in Phanerochaete chrysosporium: Activation by manganese. Journal of Bacteriology, 173(13), 4101–4106. 10.1128/jb.173.13.4101-4106.1991

33. Li, D., Alic, M., & Gold, M. H. (1994). Nitrogen regulation of lignin peroxidase gene transcription. Applied and Environmental Microbiology, 60(9), 3447–3449. 10.1128/aem.60.9.3447-3449.1994

34. Schimel, J., & Weintraub, M. (2003). The implications of exoenzyme activity on microbial carbon and nitrogen limitation in soil: A theoretical model. Soil Biology and Biochemistry, 35(4), 549–563. 10.1016/S0038-0717(03)00015-4

35. Frey, S. D., Knorr, M., Parrent, J. L., & Simpson, R. T. (2004). Chronic nitrogen enrichment affects the structure and function of the soil microbial community in temperate hardwood and pine forests. Forest Ecology and Management, 196(1), 159–171. 10.1016/j.foreco.2004.03.018

36. DeForest, J. L., Zak, D. R., Pregitzer, K. S., & Burton, A. J. (2004). Atmospheric Nitrate Deposition, Microbial Community Composition, and Enzyme Activity in Northern Hardwood Forests. Soil Science Society of America Journal, 68(1), 132–138. 10.2136/sssaj2004.1320

37. DeForest, J. L., Zak, D. R., Pregitzer, K. S., & Burton, A. J. (2005). ATMOSPHERIC NITRATE DEPOSITION AND ENHANCED DISSOLVED ORGANIC CARBON LEACHING: TEST OF A POTENTIAL MECHANISM. Soil Science Society of America Journal, 69(4), 1233–1237. 10.2136/sssaj2004.0283

38. Van Diepen, L. T. A., Frey, S. D., Sthultz, C. M., Morrison, E. W., Minocha, R., & Pringle, A. (2015). Changes in litter quality caused by simulated nitrogen deposition reinforce the N-induced suppression of litter decay. Ecosphere, 6(10), art205. 10.1890/ES15-00262.1

39. Entwistle, E. M., Romanowicz, K. J., Argiroff, W. A., Freedman, Z. B., Morris, J. J., & Zak, D. R. (2018). Anthropogenic N Deposition Alters the Composition of Expressed Class II Fungal Peroxidases. Applied and Environmental Microbiology, 84(9), e02816–17. 10.1128/AEM.02816-17

40. Freedman, Z., & Zak, D. R. (2014). Atmospheric N Deposition Increases Bacterial Laccase-Like Multicopper Oxidases: Implications for Organic Matter Decay. Applied and Environmental Microbiology, 80(14), 4460–4468. 10.1128/AEM.01224-14

41. Hesse, C. N., Mueller, R. C., Vuyisich, M., Gallegos-Graves, L. V., Gleasner, C. D., Zak, D. R., & Kuske, C. R. (2015). Forest floor community metatranscriptomes identify fungal and bacterial responses to N deposition in two maple forests. Frontiers in Microbiology, 6. 10.3389/fmicb.2015.00337

42. Aber, J., McDowell, W., Nadelhoffer, K., Magill, A., Berntson, G., Kamakea, M., McNulty, S., Currie, W., Rustad, L., & Fernandez, I. (1998). Nitrogen Saturation in Temperate Forest Ecosystems. BioScience, 48(11), 921–934. 10.2307/1313296

43. Templer, P. H., Mack, M. C., Iii, F. S. C., Christenson, L. M., Compton, J. E., Crook, H. D., Currie, W. S., Curtis, C. J., Dail, D. B., D’Antonio, C. M., Emmett, B. A., Epstein, H. E., Goodale, C. L., Gundersen, P., Hobbie, S. E., Holland, K., Hooper, D. U., Hungate, B. A., Lamontagne, S., … Zak, D. R. (2012). Sinks for nitrogen inputs in terrestrial ecosystems: A meta-analysis of ^15^ N tracer field studies. Ecology, 93(8), 1816–1829. 10.1890/11-1146.1

44. Nadelhoffer, K. J., Colman, B. P., Currie, W. S., Magill, A., & Aber, J. D. (2004). Decadal-scale fates of tracers added to oak and pine stands under ambient and elevated N inputs at the Harvard Forest (USA). Forest Ecology and Management, 196(1), 89–107. 10.1016/j.foreco.2004.03.014

45. Stevens, C. J. (2016). How long do ecosystems take to recover from atmospheric nitrogen deposition? Biological Conservation, 200, 160–167. 10.1016/j.biocon.2016.06.005

46. Zak, D. R., Freedman, Z. B., Upchurch, R. A., Steffens, M., & Kögel-Knabner, I. (2017). Anthropogenic N deposition increases soil organic matter accumulation without altering its biochemical composition. Global Change Biology, 23(2), 933–944. 10.1111/gcb.13480

47. Gurmesa, G. A., Wang, A., Li, S., Peng, S., De Vries, W., Gundersen, P., Ciais, P., Phillips, O. L., Hobbie, E. A., Zhu, W., Nadelhoffer, K., Xi, Y., Bai, E., Sun, T., Chen, D., Zhou, W., Zhang, Y., Guo, Y., Zhu, J., … Fang, Y. (2022). Retention of deposited ammonium and nitrate and its impact on the global forest carbon sink. Nature Communications, 13(1), 880. 10.1038/s41467-022-28345-1

48. Shao, P., Zeng, X., Moore, D. J. P., & Zeng, X. (2013). Soil microbial respiration from observations and Earth System Models. Environmental Research Letters, 8(3), 034034. 10.1088/1748-9326/8/3/034034

49. Carvalhais, N., Forkel, M., Khomik, M., Bellarby, J., Jung, M., Migliavacca, M., Μu, M., Saatchi, S., Santoro, M., Thurner, M., Weber, U., Ahrens, B., Beer, C., Cescatti, A., Randerson, J. T., &., SaatchiReichstein, M. (2014). Global covariation of carbon turnover times with climate in terrestrial ecosystems. Nature, 514(7521), 213–217. 10.1038/nature13731

50. Peñuelas, J., Ciais, P., Canadell, J. G., Janssens, I. A., Fernández-Martínez, M., Carnicer, J., Obersteiner, M., Piao, S., Vautard, R., & Sardans, J. (2017). Shifting from a fertilization-dominated to a warming-dominated period. Nature Ecology & Evolution, 1(10), 1438–1445. 10.1038/s41559-017-0274-8

51. Hartley, I. P., Hill, T. C., Chadburn, S. E., & Hugelius, G. (2021). Temperature effects on carbon storage are controlled by soil stabilisation capacities. Nature Communications, 12(1), 6713. 10.1038/s41467-021-27101-1

52. Fernández-Martínez, M., Peñuelas, J., Chevallier, F., Ciais, P., Obersteiner, M., Rödenbeck, C., Sardans, J., Vicca, S., Yang, H., Sitch, S., Friedlingstein, P., Arora, V. K., Goll, D. S., Jain, A. K., Lombardozzi, D. L., McGuire, P. C., & Janssens, I. A. (2023). Diagnosing destabilization risk in global land carbon sinks. Nature, 615(7954), 848–853. 10.1038/s41586-023-05725-1

53. Steidinger, B. S., Crowther, T. W., Liang, J., Van Nuland, M. E., Werner, G. D. A., Reich, P. B., Nabuurs, G. J., de-Miguel, S., Zhou, M., Picard, N., Herault, B., Zhao, X., Zhang, C., Routh, D., & Peay, K. G. (2019). Climatic controls of decomposition drive the global biogeography of forest-tree symbioses. Nature, 569(7756), 404–408. 10.1038/s41586-019-1128-0

54. Litzow, M. A., & Hunsicker, M. E. (2016). Early warning signals, nonlinearity, and signs of hysteresis in real ecosystems. Ecosphere, 7(12). 10.1002/ecs2.1614

55. Yost, J. L., & Hartemink, A. E. (2019). Soil organic carbon in sandy soils: A review In Advances in Agronomy (Vol. 158, pp. 217–310). Elsevier. 10.1016/bs.agron.2019.07.004

56. Wu, J., Zhang, H., Pan, Y., Cheng, X., Zhang, K., & Liu, G. (2023). Particulate organic carbon is more sensitive to nitrogen addition than mineral-associated organic carbon: A meta-analysis. Soil and Tillage Research, 232, 105770. 10.1016/j.still.2023.105770

57. Burton, A. J., Ramm, C. W., Pregitzer, K. S., & Reed, D. D. (1991). Use of multivariate methods in forest research site selection. Canadian Journal of Forest Research, 21(11), 1573–1580. 10.1139/x91-219

58. Franzluebbers, A. J., Haney, R. L., Hons, F. M., & Zuberer, D. A. (1996). Determination of Microbial Biomass and Nitrogen Mineralization following Rewetting of Dried Soil. Soil Science Society of America Journal, 60(4), 1133–1139. 10.2136/sssaj1996.03615995006000040025x

59. Franzluebbers, A. J., Haney, R. L., Honeycutt, C. W., Schomberg, H. H., & Hons, F. M. (2000). Flush of Carbon Dioxide Following Rewetting of Dried Soil Relates to Active Organic Pools. Soil Science Society of America Journal, 64(2), 613–623. 10.2136/sssaj2000.642613x

60. Wade, J., Culman, S. W., Hurisso, T. T., Miller, R. O., Baker, L., & Horwath, W. R. (2018). Sources of Variability that Compromise Mineralizable Carbon as a Soil Health Indicator. Soil Science Society of America Journal, 82(1), 243–252. 10.2136/sssaj2017.03.0105

61. Franzluebbers, A. J. (2016). Should Soil Testing Services Measure Soil Biological Activity? Agricultural & Environmental Letters, 1(1), 150009. 10.2134/ael2015.11.0009

62. Saiya-Cork, K. R., Sinsabaugh, R. L., & Zak, D. R. (2002). The effects of long term nitrogen deposition on extracellular enzyme activity in an Acer saccharum forest soil. Soil Biology and Biochemistry, 34(9), 1309–1315. 10.1016/S0038-0717(02)00074-3

63. Freedman, Z. B., Romanowicz, K. J., Upchurch, R. A., & Zak, D. R. (2015). Differential responses of total and active soil microbial communities to long-term experimental N deposition. Soil Biology and Biochemistry, 90, 275–282. 10.1016/j.soilbio.2015.08.014

64. Freedman, Z. B., Upchurch, R. A., Zak, D. R., & Cline, L. C. (2016). Anthropogenic N Deposition Slows Decay by Favoring Bacterial Metabolism: Insights from Metagenomic Analyses. Frontiers in Microbiology, 7. 10.3389/fmicb.2016.00259

65. Ben-Shachar, M., Lüdecke, D., & Makowski, D. (2020). effectsize: Estimation of Effect Size Indices and Standardized Parameters. Journal of Open Source Software, 5(56), 2815. 10.21105/joss.02815

66. Cohen, J. (1988). *Statistical power analysis for the behavioral sciences* (2nd ed). L. Erlbaum Associates.

67. Kassambara A (2023). rstatix: Pipe-Friendly Framework for Basic Statistical Tests. R package version 0.7.2, https://rpkgs.datanovia.com/rstatix/.

68. R Core Team (2022). R: A Language and Environment for Statistical Computing. R Foundation for Statistical Computing, Vienna, Austria. https://www.R-project.org/.

69. RStudio Team (2022). RStudio: Integrated development environment for R. Boston, MA. http://www.rstudio.com/.

